# Predictive gaze orienting during navigation in virtual reality

**DOI:** 10.1101/2025.09.24.678200

**Authors:** Vasiliki Kondyli, Marcin Leszczyński

## Abstract

Natural vision is an active, predictive process guided by expectations about when and where information will appear. Yet how gaze is shaped in dynamic, multisensory environments remains poorly understood. Using immersive virtual reality with eye-tracking, we examined oculomotor behavior during naturalistic navigation. Participants cycled through a virtual city while avatar cyclists, first heard as overtaking them from behind via spatialized auditory cues, later became visible as they passed. Auditory cues triggered anticipatory gaze shifts to expected locations, indicating that eye movements were guided by auditory predictions rather than reactive visual responses. Violations of auditory–spatial expectations elicited longer fixations. Critically, removing auditory cues impaired predictive gaze orienting, delayed gaze orienting and increased collisions with obstacles. These findings demonstrate that auditory input fundamentally shapes predictive models guiding visual exploration and adaptive behavior in dynamic environments, underscoring the multisensory basis of active perception in real-world interactions.

## Introduction

Imagine cycling through a narrow city street during rush hour. Other vehicles approach head-on while some overtake from behind. Glancing over your shoulder isn’t always possible – or necessary. Instead, you rely on auditory cues to anticipate when and where someone might pass. Here, we explored how these sounds are used to orient visual exploration and support adaptive behavior. Perception in natural environments is an active, predictive, and multisensory process. Most sensory inputs are acquired through active motor sampling routines (Ahissar & Arieli, 2001; Kleinfeld et al., 2006; Schroeder et al., 2010; Leszczynski & Schroeder, 2019; Leszczynski et al., 2021; Nentwich et al., 2023; Leszczynski et al., 2025). In vision, saccadic eye movements, head turns, and body shifts continuously reorient the fovea to task-relevant locations (Yarbus, 1967; Gilchrist et al., 1997). A central question in vision science is how the brain guides these movements during real-world behaviour. Accumulating evidence suggests that gaze is guided by predictive models that integrate prior knowledge and contextual information to anticipate where and when relevant stimuli will appear, Henderson, 2003, 2017; Brockmole & Henderson, 2006; Tatler et al., 2011; Hayhoe et al. 2012; Goettker et al. 2021; Boettcher et al. 2022).

Spatial predictions have been shown to shape visual exploration: objects in expected locations are detected more efficiently, while unexpected placements trigger prolonged and repeated fixations (Võ et al., 2009; Henderson et al., 2007). Temporal predictions have also emerged as key to visual behaviour in dynamic contexts. Skilled as well as naive observers direct their gaze toward predicted event locations before the stimulus of interest appears there, revealing anticipatory sampling of the visual scene (Land & McLeod, 2000; Diaz et al., 2013; Goettker et al. 2021). These findings suggest that active vision is guided based on internal models of the world that not only contain predictions about *where*, but also *when* relevant events will occur. Predictive gaze orienting is also fundamental for adaptive behaviour in dynamic environments. It enables timely detection of potential hazards, thereby supporting rapid and appropriate behavioural responses (Diaz et al., 2013; Hayhoe & Ballard, 2014; Bakst & McGuire, 2021). Failures in using predictive cues can lead to delayed reactions and missed detections, increasing the risk of collisions or falls during navigation and driving (Crundall et al., 1999; Underwood et al., 2002; Wolfe et al., 2017; Tuhkanen et al., 2019, 2021; Vansteenkiste et al. 2013). Recent models emphasize that active vision relies on continuously updated internal models that integrate multisensory cues to optimize both perception and action (Henderson, 2017; Leszczynski & Schroeder, 2019; Shalev et al., 2024).

Despite these advances, it remains unknown how multisensory cues contribute to these spatiotemporal predictions in dynamic, naturalistic environments. Most prior work has focused on static, unisensory scenes. However, real-world perception often unfolds in rapidly changing environments where vision is incomplete and often supplemented by sound. Auditory cues, in particular, can signal the presence and trajectory of objects outside the current field of view, providing crucial predictive information for guiding eye movements. Despite growing evidence for audio-visual interaction during active vision (Rummukainen & Mendonça, 2016; Leszczynski et al., 2023; Gehmacher et al., 2024), how auditory cues shape oculomotor behaviour in naturalistic contexts—and whether they enhance real-world adaptive behaviour—remains poorly understood. To address this, we used immersive virtual reality (VR) with continuous eye-tracking to investigate how auditory information influences predictive gaze behaviour during naturalistic navigation in VR. Participants navigated a virtual city on bicycles while occasionally being overtaken by audio-visual avatars (bikes or motorbikes). Mirroring real-life experiences, these overtaking events were preceded by directionally localized sounds signaling the avatars’ approach. Participants first heard an avatar cyclist coming from behind, the sound growing closer, before the avatar visually overtook them from either the left or right side. We tested three hypotheses: (1) auditory cues guide predictive gaze shifts toward expected avatar locations even before it becomes visible, (2) violations of spatiotemporal predictions lead to increased visual scrutiny with prolonged visual exploration of violating objects, and (3) auditory-driven predictions and associated predictive gaze orienting facilitates adaptive responses to environmental hazards.

Our findings show that gaze was predictively directed toward expected avatar locations up to 240 ms before the visual onset, revealing an influence of auditory-based prediction on gaze orienting. When avatars violated these predictions, gaze patterns reflected increased exploration, with fixations concentrated at the anticipated—but incorrect—location. Notably, 51.5% of first saccades following the avatar’s entry into the visual field were directed toward the predicted location, even when the avatar appeared on the opposite side. Finally, disruption of these auditory cues impaired predictive oculomotor orienting and increased collision rates with virtual obstacles, confirming a behavioural advantage conferred by predictions in active sensing. Together, these results demonstrate that auditory cues continuously update internal models that guide active visual exploration and support adaptive behaviour in complex, dynamic environments.

## Results

### Predictive gaze shifts toward anticipated locations of approaching audio-visual objects

Participants cycled along a 25-kilometer virtual city route in first-person view, following directional wayfinding signs toward the city of “Liberty” to reach a specified destination (Figure 1A). Avatars of cyclists and motorcyclists overtook them at randomly selected intervals (mean = 28 s; range: 1.5–55 s), mimicking real-world urban navigation. The sound of the cyclist included the rhythmic whirring of the chain, while the motorcyclist’s sound featured a low-frequency engine rumble. The setup accounted for natural sound propagation and environmental interactions to replicate real-world acoustic conditions, including changes in intensity with distance and the Doppler effect. The avatars first became detectable through auditory cues—originating from beyond the visual field—and only later appeared visually as they passed. On average, participants heard each approaching avatar for 6 seconds before it entered their field of view (range: 2.6–19.1 s), depending on head rotation and cycling velocity. Over the course of the task (average of 21 minutes), each participant experienced 40 overtakes (20 from the left, 20 from the right, randomly ordered). Continuous eye tracking (50 Hz) revealed systematic gaze shifts oriented toward the direction of the impending avatar, beginning even before the avatar’s visual onset.

**Figure 1.**
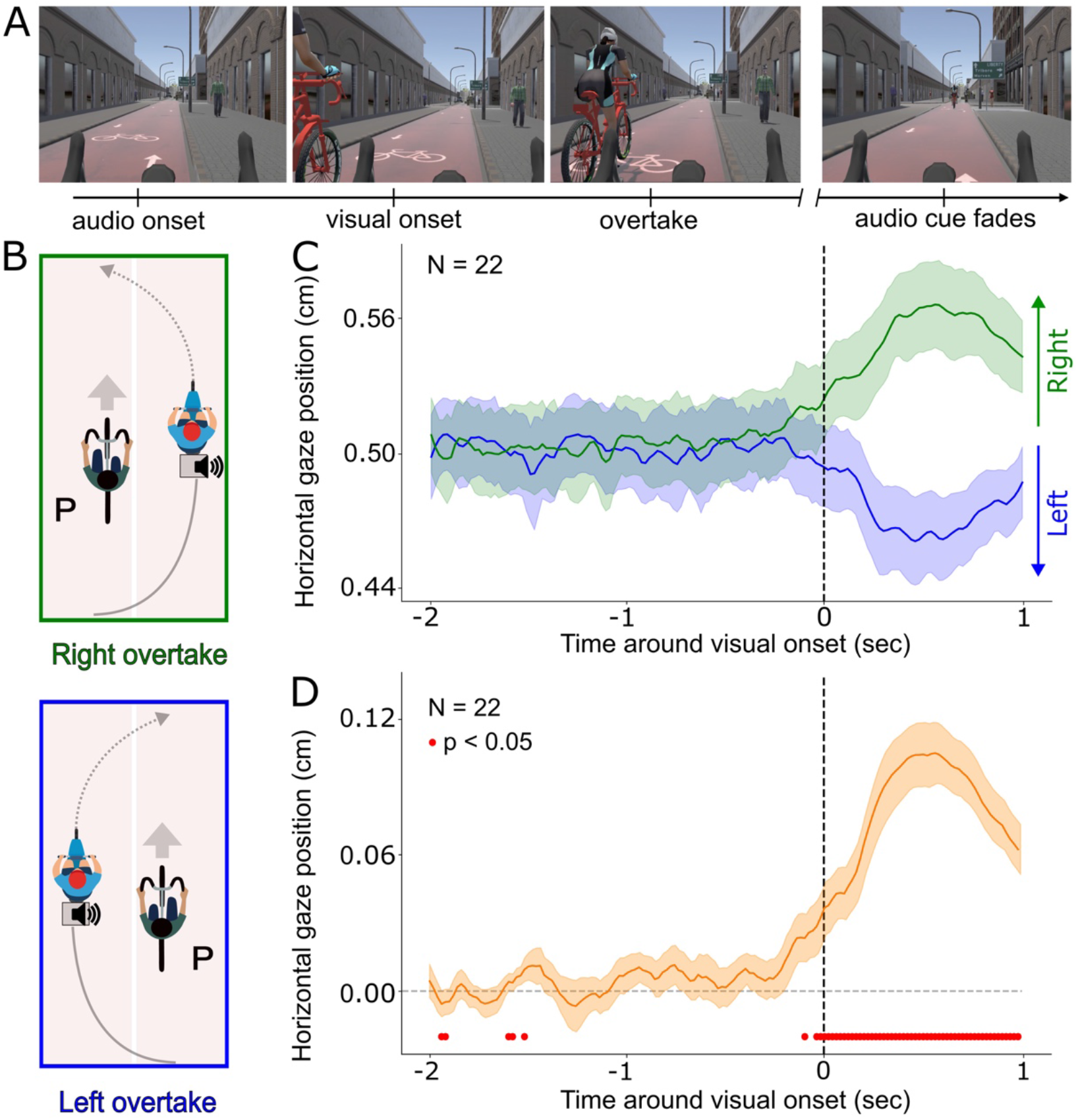
Orienting of eye movements in multisensory environment (Experiment 1). **(A)** Participants navigated a virtual city from a first-person cycling perspective. The panels illustrate a representative overtaking event: an approaching avatar is initially audible but not visible (far left), enters the visual field from behind (second panel), overtakes the participant (third panel), and then moves ahead (far right). This sequence mirrors real-world sensory dynamics, in which auditory cues precede and guide visual orienting toward task-relevant events. **(B)** Spatial configuration of overtaking events, with avatars approaching and passing the participant’s bicycle from either the left or right side. This design allowed us to systematically examine lateralized auditory cues and their influence on anticipatory gaze shifts prior to visual appearance of the avatar. The solid line represents the trajectory of the audio source, while the dashed line represents the trajectory of the visual avatar. **(C)** Horizontal gaze position during left (blue) and right (green) overtakes. **(D)** Towardness index quantifies horizontal gaze orientation toward the auditory-cued location, with positive values reflecting shifts in the direction of the approaching avatar. Red dots along the x-axis denote time points where towardness index significantly diverged from baseline (p < 0.05; Benjamini-Hochberg corrected; N = 22). In panels C and D, time is aligned to avatar visual onset (0 s; vertical dashed line), and the y-axis indicates horizontal gaze position. Shaded regions represent SEM.

To investigate the dynamics of auditory-guided gaze orienting, we aligned gaze trajectories to the moment each avatar entered the visual field of the observer (i.e., visual onset). This analysis focused on eye movements; head movements were not analyzed. Separate analyses for left and right overtakes revealed anticipatory horizontal deviations in gaze toward the auditory cue (Figure 1B). To quantify gaze orienting toward auditory-cued locations, we computed a unified “towardness” index by combining gaze positions during leftward and rightward overtakes, capturing displacement toward the cued location irrespective of side (Figure 1C; for similar approach see Draschkow et al., 2022). Relative to a baseline window (−2 to −1.5 s), towardness values showed a significant increase beginning at 100 ms before visual onset (Wilcoxon signed-rank tests, all p < 0.05, all ∣z∣ > 2.32, N = 22; Benjamini-Hochberg FDR corrected). These anticipatory gaze shifts emerged prior to visual appearance of the avatar, demonstrating that gaze was relocated towards the location where the avatar was expected to appear— driven by auditory cues rather than triggered by visual onset. These transient shifts were detectable as early as 1.5 seconds before visual onset, followed by a later increase 100 ms before visual onset. These data suggest that participants continuously update their spatial expectation, predictively (i.e., before an object appears there), directing their gaze and guiding active vision based on auditory input.

These findings show that, even without explicit instructions to attend to overtaking avatars, participants’ gaze was systematically directed toward the anticipated point of visual entry. Gaze shifts emerged shortly before the avatar appeared, indicating that auditory cues – informing about objects outside the visual field – trigger anticipatory visual orienting. This observed behaviour suggests that the auditory system contributes to active visual exploration by integrating spatiotemporal information about objects outside of the field of view. Such anticipatory gaze shifts likely optimize foveal access to behaviourally relevant stimuli, supporting efficient visuomotor coordination in dynamic, multisensory environments. The results demonstrate that participants use multisensory cues to continuously construct and update internal models of the environment to predictively allocate gaze in service of adaptive perception and action.

### Increased visual exploration of objects violating predictions

In Experiment 1, we found that participants used auditory cues from out-of-view avatars to shift their gaze toward the expected location of visual appearance. Building on prior work demonstrating that violations of audio-spatial regularities enhance visual scrutiny (Henderson et al., 1999; Võ et al., 2009), we hypothesized that prediction violations in dynamic, multisensory settings would similarly intensify visual exploration. Specifically, we expected prolonged duration of fixation at the visual avatar when it appears in unexpected locations. To test this, we conducted Experiment 2 with a new cohort of participants using a task matched to Experiment 1 but incorporating systematic violations of spatiotemporal predictions. Participants navigated a virtual city (Figure 2A–B), where avatars of cyclists and motorcyclists overtook them at variable intervals. On 70% of trials (congruent condition), auditory cues correctly predicted the side from which the avatar would appear. On 30% of trials (incongruent condition), visual avatars appeared on the side opposite to that indicated by the auditory cue—violating spatial, but not temporal, expectations.

**Figure 2.**
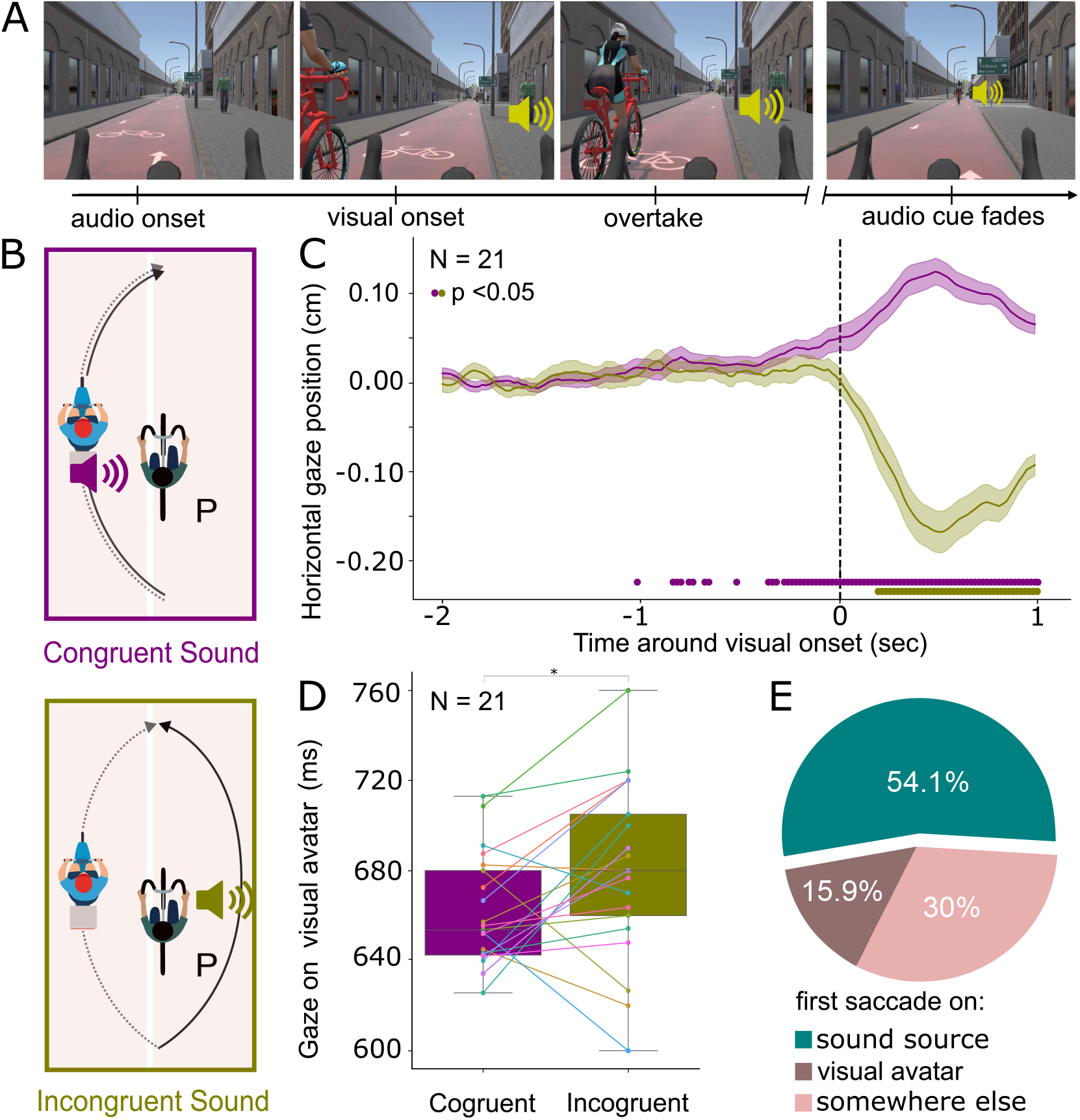
The influence of prediction violation on gaze orienting (Experiment 2). **(A)** Participants navigated a virtual city from a first-person cycling perspective. The panels illustrate a representative overtaking event with an incongruent sound: an approaching avatar is initially audible but not visible (far left), enters the visual field from the left side while its audio source mirrors the path from the right side (second panel), the avatar and the audio source overtake the participant (third panel), and then moves ahead (far right). **(B)** Schematic representation of avatar trajectories in two conditions. In congruent trials (top; 70 %), the avatar is heard approaching from one side (e.g., left) and appears visually on the same side. In incongruent trials (bottom; 30 % of trials), the avatar is heard from one side but appears on the opposite side. The solid line represents the trajectory of the audio source, while the dashed line represents the trajectory of the visual avatar. **(C)** Gaze orienting is quantified using a towardness index, where positive values reflect gaze shifts toward the auditory-cued side. Time series show mean gaze dynamics for congruent (purple) and incongruent (green) trials, aligned to visual onset (dashed vertical line). Dots along the x-axis indicate significant departure from baseline (Wilcoxon signed-rank test, p < 0.05, FDR-controlled for multiple comparisons using the Benjamini-Hochberg procedure). Shaded areas denote ±SEM. **(D)** Boxplots show dwell time on the visual avatar following its visual appearance, across conditions. Central lines indicate medians; boxes denote interquartile ranges; whiskers extend to non-outlier extremes. Asterisks indicate p < 0.05 (Wilcoxon signed-rank test). **(E)** Proportion of incongruent trials in which the first saccade after the visual onset of the avatar was directed toward the expected (i.e., auditory-cued or the sound source) location, the actual visual location of the avatar, or other parts of the environment.

We first asked whether gaze behaviour replicated the anticipatory orienting observed in Experiment 1. In congruent trials, participants reliably oriented their gaze toward the cued location as early as 360 ms prior to the avatar’s visual onset (Wilcoxon signed-rank test; all p < 0.05, all ∣z∣ > 2.28; controlled for multiple comparisons using the Benjamini-Hochberg procedure; N = 21; Figure 2C), closely replicating the temporal dynamics of Experiment 1. In incongruent trials, gaze shifted significantly only after the visual onset, becoming evident 200 ms after the avatar’s visual appearance (i.e. after the visual onset) (Wilcoxon signed-rank test; all p < 0.05, all ∣z∣ > 2.76; controlled for multiple comparisons using the Benjamini-Hochberg procedure; N = 21). In the incongruent condition, similarly to the congruent condition, the gaze was initially directed toward the auditory-cued location, reflecting the influence of the auditory cue on gaze orienting (Figure 2C). However, following the onset of the visual avatar, observers adjusted their gaze to align with the unexpected new location of the visual avatar, overriding the earlier gaze behaviour driven by the auditory cue. While early gaze patterns in the incongruent condition initially resembled those observed in the congruent condition, this effect did not reach statistical significance after correction for multiple comparisons—likely due to increased variance introduced by the corrective saccades made around the time of the visual appearance of the visual avatar (see also Figure).

Next, we examined whether violations of audio-spatial predictions modulated visual exploration following stimulus onset. Consistent with our hypothesis, dwell time on the avatar was longer in the incongruent condition compared to the congruent condition (M = 680 ms, SD = 38.5 vs. M = 660 ms, SD = 24.6; Wilcoxon signed-rank test; z = 2.31, p = 0.019; N = 21; Figure 2D). This suggests that predictions’ violations elicited prolonged fixations, possibly reflecting the need to identify the unexpected stimuli, resolve the conflict between prior expectations and incoming sensory input and integrate the unexpected stimulus with the rest of the scene.

Importantly, in 54.1% of incongruent trials, the first saccade following the avatar’s visual onset was directed toward the anticipated (i.e., auditory-cued) – but incorrect – side of the screen (Figure 2E), despite the onset of a salient visual object on the opposite side. Together, these results provide evidence that multisensory expectations shape anticipatory gaze behaviour in dynamic environments. Auditory cues establish spatiotemporal predictions that guide visual exploration. Violating these predictions results in longer fixations on these violating objects, which likely reflects efforts to reconcile mismatches between prediction and perception.

### Predictions support adaptive behaviour

Across Experiments 1 and 2, we showed that during dynamic multisensory events, participants sampled information across modalities to form predictions about object trajectories. Specifically, auditory cues enabled the generation of spatiotemporal predictions for objects outside the visual field, with violations of these predictions leading to enhanced visual inspection.

To test whether such predictions facilitate adaptive behaviour, we conducted Experiment 3 with a new group of participants (N = 20). Participants navigated a virtual city (Figure 3A) where avatars of cyclists and motorcyclists overtook them at randomized intervals, consistent with the timing parameters of Experiments 1 and 2. Some avatars (in 60% of all trials) carried a package, and in half of these trials (30% of all trials), the package was dropped in front of the participant following the overtaking manoeuvre, requiring an immediate behavioural response to avoid collision. On average, the box was dropped 500-800 ms after becoming visible. We introduced two experimental conditions: one with auditory cues (audio-on, enabling spatiotemporal predictions) and one without (audio-off). All parameters were identical across conditions except for the availability of auditory information.

**Figure 3.**
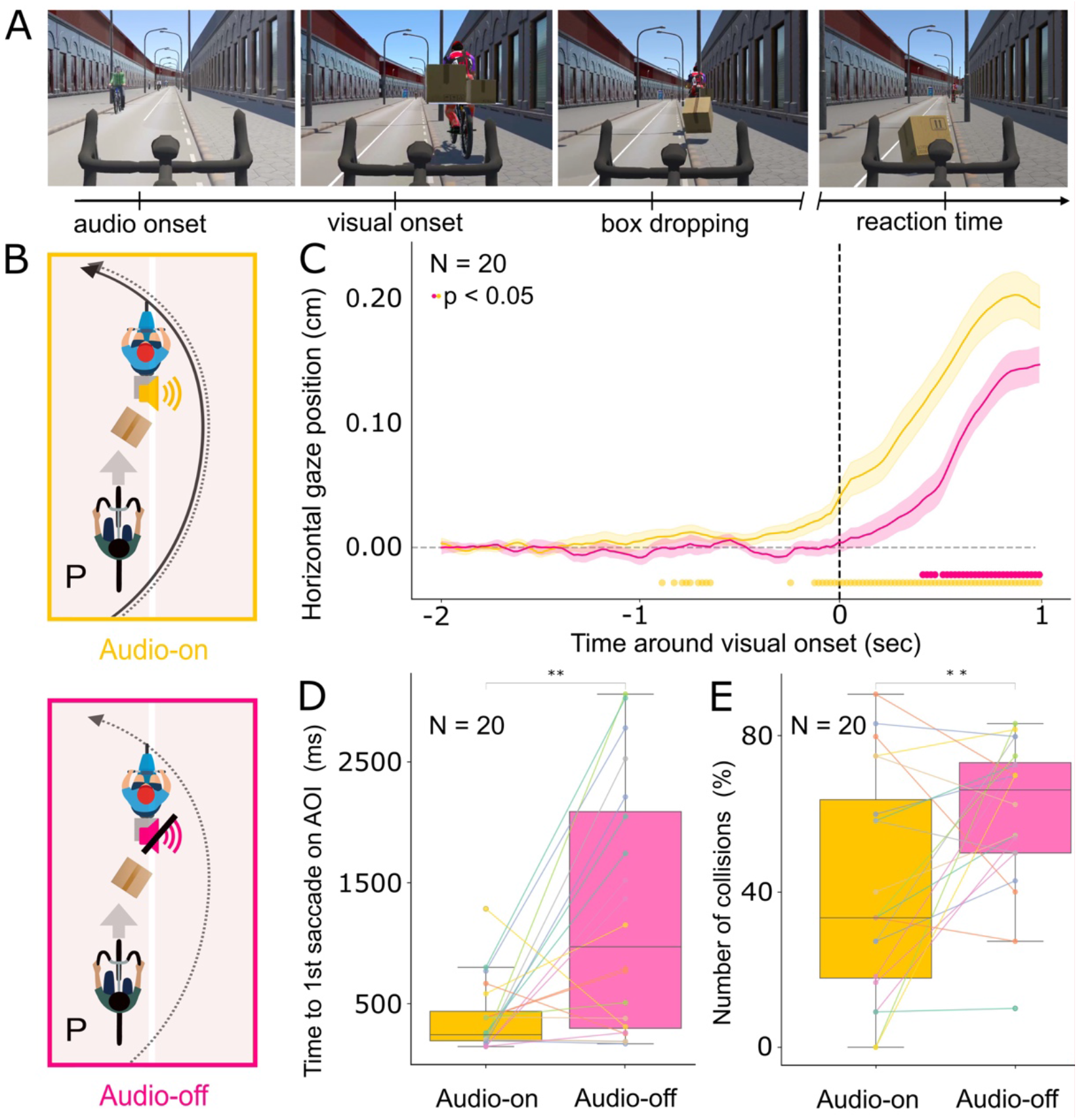
Predictive auditory cues enhance adaptive visuomotor responses during dynamic events (Experiment 3). **(A)** Participants navigated a virtual city from a first-person cycling perspective. The panels illustrate a representative overtaking event involving a dynamic obstacle: an approaching avatar is initially heard (audio-on condition only; far left), enters the visual field from behind and overtakes the participant (second panel), and subsequently drops a box directly into the participant’s path (30 % of all trials; third panel), necessitating an immediate steering response to avoid collision (far right). **(B)** Schematic illustration of the virtual environment and the relative positions of the cyclist and avatar at the time of the box drop, across two experimental conditions. In the audio-on condition (top), auditory cues provide spatially predictive information about the overtaking avatar’s trajectory, as in Experiments 1–2. In the audio-off condition (bottom), no auditory cues are available throughout the session, eliminating the possibility of generating auditory-based prediction. The solid line represents the trajectory of the audio source, while the dashed line represents the trajectory of the visual avatar. **(C)** Time course of the towardness index, quantifying horizontal gaze orientation toward the side of the approaching avatar. Positive values indicate gaze shifts in the direction of the overtaking side. Colored dots along the x-axis indicate time points where gaze significantly departed from baseline (yellow: audio-on; pink: audio-off; p < 0.05, Wilcoxon signed-rank test, controlled for multiple comparisons with Benjamini-Hochberg procedure; N = 20). Time (x-axis) is aligned to the visual onset of the overtaking avatar (0 s; dashed vertical line). Shaded regions indicate ±SEM. **(D)** Latency of the first saccade to the area of interest (includes the box and the avatar) after it was dropped shown separately for each condition. **(E)** Proportion of trials in which participants collided with the dropped box, comparing predictive (audio-on) and non-predictive (audio-off) conditions. Box plots in D and E represent the 25^th^, median, and 75^th^ percentiles, with whiskers extending to extreme values. Double asterisks indicate significant differences (p < 0.01, Wilcoxon signed-rank test).

Consistent with earlier findings, in the predictive condition (i.e., audio-on), participants oriented their gaze toward the anticipated location of the approaching avatar before its visual appearance in the visual field of the observer. We identify two significant time windows of anticipatory gaze shifts: an early window 660–900 ms before visual onset, and a second window beginning just 80 ms prior to the onset (Wilcoxon signed-rank test, all p < 0.05; all ∣z∣ > 2.28; controlled for multiple comparisons with Benjamini-Hochberg procedure; N = 20; Figure 3C). In contrast, in the condition without predictions (i.e., audio-off), anticipatory gaze orienting was absent; participants shifted gaze toward the avatar only 400 ms after its visual appearance (Wilcoxon signed-rank test, all p < 0.05; all ∣z∣ > 2.69; N = 20; controlled for multiple comparisons with Benjamini-Hochberg procedure) consistent with previous studies on reflective gaze orienting towards a visual stimulus popping out in the peripheral view (Vencato, et al., 2020).To better understand how the availability of auditory cues influences gaze orienting, we examined the time to first saccade to the area of interest (AOI; including the visual avatar and the box) following the box dropped in the audio-on and audio-off conditions. Saccades were significantly faster in the audio-on compared to the audio-off condition (M = 391 ms, SD = 296.72 vs. M = 1265 ms, SD = 1034.39 ms; Wilcoxon signed-rank test: z = −2.80, p = 0.005; N = 20; Figure 3D). We also analyzed dwell time during the interval when both the agent and the box were visible but prior to the box dropping, focusing only on trials where the box dropped. Within this 500–800 milliseconds window, there was no significant difference in dwell time on the AOIs between the audio-on and audio-off conditions (321 milliseconds vs. 371 milliseconds, respectively; Wilcoxon signed-rank test: z = −1.49, p = 0.135; N = 20).

Finally, predictive auditory cues substantially reduced collision rates: participants collided with obstacles less frequently in the audio-on condition compared to the audio-off condition (41.42% vs. 60.06%; Wilcoxon signed-rank test: z = 2.35, p = 0.017; N = 20; Figure 3E). Together, these findings demonstrate that spatiotemporal predictions generated based on auditory cues not only guide anticipatory gaze but also enhance adaptive behaviour in dynamic, multisensory environments.

## Discussion

Using virtual reality combined with simultaneous eye-tracking, we investigated how gaze is guided during navigation through a dynamic audiovisual environment. While cycling through a virtual city, participants were occasionally overtaken by audiovisual avatars of other cyclists, allowing us to examine gaze shifts elicited by these overtaking events. Our findings reveal three key insights. First, auditory information plays a crucial role in directing gaze toward locations where audible objects are expected to enter the visual field. These gaze shifts occurred before the avatar became visible, indicating they were driven by auditory-based predictions about the avatar’s trajectory rather than reactive, visually driven responses. Second, objects that violated these predictions—avatars emerging from the opposite side of the expected location—elicited prolonged and more frequent fixations, consistent with enhanced exploration of unexpected stimuli. Third, auditory predictions were critical for adaptive behaviour— removing access to auditory-based predictions (while keeping visual representation of avatars comparable) impaired participants’ ability to react to road conditions, leading to a higher rate of collisions. Together, our findings support a model in which auditory cues continuously inform predictive models that guide gaze toward locations where relevant changes are anticipated. This predictive mechanism optimizes visual exploration and enhances adaptive behaviour in dynamic, multisensory environments.

### The predictive gaze control in dynamic multisensory environments

Understanding how humans allocate gaze in real-world contexts has long been a central question in cognitive science and vision science. Two major theoretical frameworks have dominated the field. One posits that gaze is largely governed by bottom-up features—such as contrast, color, and luminance— captured by early saliency models (Itti & Koch, 2001; Parkhurst et al., 2002; Borji et al., 2013). While recent advances in deep learning have enhanced the predictive accuracy of these models (Kümmerer et al., 2017), they remain limited in their capacity to account for the temporal evolution of gaze or the integration of non-visual cues. In contrast, a growing body of work emphasizes the role of top-down mechanisms—semantic relevance, prior knowledge, and task demands—in shaping gaze behaviour (Henderson, 2020; Henderson et al., 2009; Nyström & Holmqvist, 2008; Onat et al., 2014). According to this view, gaze is deployed not merely in response to what is visually salient, but rather to what is informationally valuable based on contextual (Land & McLeod, 2000; Hwang et al., 2011; Hayhoe et al. 2012; Diaz et al., 2013; Võ et al., 2019; Goettker et al, 2021; Peacock et al., 2019; 2023; Pedziwiatr et al., 2023) or memory derived predictions (Bar, 2009; Diaz et al., 2013; Gorman et al., 2012).

Our findings demonstrate that in dynamic, multisensory environments, gaze is predictively oriented by auditory cues. Participants consistently oriented their gaze toward the expected location of upcoming visual events, guided by spatial information in sound. Strikingly, in Experiment 2, over half of initial saccades landed on the predicted side even when the visual stimulus appeared elsewhere—revealing that cross-modal predictions can override incoming visual input. Removing auditory cues in Experiment 3 delayed gaze shifts by over a second and increased hazard collisions, indicating clear behavioural costs of disrupted prediction. These results support a model in which gaze is proactively guided by internal, multisensory models rather than driven solely by visual input.

In line with predictive processing theories (Rao & Ballard, 1999; Clark, 2013; Rao, 2024), our data show that auditory cues shape visual sampling (see also McDonald et al., 2000). Auditory signals dynamically influence internal models, facilitating rapid, goal-directed gaze shifts that support adaptive behaviour. These findings extend prior work on anticipatory vision in natural tasks (Jovancevic-Misic & Hayhoe, 2009; M. Land & Tatler, 2009; Hayhoe, 2017; Tong et al., 2017) by demonstrating that cross-modal predictions are not only pervasive but functionally essential for safe and efficient interaction with complex multisensory environments.

### The role of the auditory system in guiding active visual exploration

While the auditory system is often viewed as a detector of distal events, a growing body of evidence suggests that one of its core functions is also to direct visual attention toward behaviourally relevant locations (Heffner & Heffner, 1992, 2007; Spence & Driver, 1997; McDonald et al., 2000; Kubovy & Van Valkenburg, 2001; Heffner et al., 2001; Ho & Spence 2005; Kean, & Crawford, 2008). Comparative studies show that species with a broad field of best view, such as those with a visual streak, require less precise auditory localization than species with a narrow, foveated field of best vision (Heffner & Heffner, 1992). The authors proposed that the need to direct the area of highest visual acuity toward a sound source for further examination plays a critical role in shaping sound localization abilities across species (Heffner & Heffner, 1992). This view is also supported by proposals that the dorsal auditory stream acts as a vision-guiding system, linking auditory spatial maps with visuomotor circuitry (Arnott & Alain, 2011; Rauschecker, 2011).

Consistent with this model, previous evidence showed that auditory cues enhance allocation of covert spatial attention (Spence & Driver, 1997; McDonald et al., 2000; Ahveninen et al., 2019; Hülsdünker et al., 2021). Our findings further reveal that auditory cues orchestrate not only covert attention but also overt gaze behaviour. In naturalistic, multisensory settings, participants used auditory cues to anticipate where and when visual events would occur, shifting gaze toward predicted locations even before any visual input was available. These anticipatory saccades were not only robust but behaviourally relevant: when auditory information was withheld, gaze orienting was delayed, and participants were more likely to collide with sudden road hazards. This indicates that auditory cues instantiate spatiotemporal predictions that prepare the visual system for targeted sampling and rapid response. Rather than passively responding to visual salience, gaze is proactively guided by cross-modal expectations— underscoring the predictive, integrative nature of real-world perception and action.

### Predictive gaze orienting supports adaptive behaviour

Adaptive behaviour requires timely sampling and integration of sensory information. Whether walking across uneven terrain (Matthis et al., 2018), steering a vehicle (Land & Lee, 1994; Land & Tatler, 2001) or making economic choices (Armel et al., 2008; Krajbich et al., 2010), where we look—and when— directly influences our decisions and actions. Gaze acts not only as a window into internal cognitive states but as a proactive mechanism for reducing uncertainty, anticipating future events, and guiding behaviour in complex, temporally unfolding contexts (Hayhoe & Ballard, 2005; Land & Tatler, 2009; Tatler et al., 2011). In naturalistic environments, the strategic allocation of gaze allows organisms to sample the most behaviourally relevant information at the right moment, facilitating rapid and efficient responses (Kircher et al., 2022; Kerautret et al. 2023; Keshava et al. 2024).

Our findings show that timely gaze orienting towards the predicted location is critical for adaptive behaviour. Eliminating auditory cues delayed orienting and increased road collisions, showing that auditory cues and associated predictive orienting help to rapidly respond to road hazards, such as avoiding a box dropped from an overtaking bicycle. Our results also help understand why listening to music or talking on a phone, as well as the quietness of electric cars at low speeds, all negatively influence auditory perception of road users and their overall performance and increase risks of a crash (Stelling-Kończak et al., 2015; Springer-Teumer et al., 2023).

### Exploring objects that violate predictions

Natural environments are governed by statistical regularities—spatial and semantic patterns that the brain exploits to reduce uncertainty and guide behaviour. In static scenes, a learned set of expectations about the typical spatial arrangement and meaning of objects facilitates perception and goal-directed attention (Võ & Henderson, 2009; Võ, 2021). Violating these expectations – by placing objects in unusual locations (e.g., a toaster on the floor) or introducing semantically inconsistent elements (e.g., an octopus in a backyard) – impairs gaze orienting and associated attention allocation as evidenced by longer fixations and increased search times (Loftus & Mackworth, 1978; Henderson et al., 2009; Öhlschläger & Võ, 2017).

Our study extends these findings into dynamic, multisensory environments by showing that violations of spatiotemporal predictions—driven by auditory cues—shape visual exploration. We observed that avatars violating auditory-based expectations were fixated for longer durations, consistent with prior findings that prediction errors attract attention (Võ & Henderson, 2009; Horstmann & Herwig, 2015; Tang et al., 2018). Furthermore, the initial fixation on the avatar was delayed in incongruent conditions compared to congruent ones. Importantly, this prolonged attention was associated with a behavioural cost: participants detected incongruent avatars more slowly than congruent ones, suggesting that prediction violations delay perceptual processing. These results highlight the role of auditory cues in shaping internal models that guide visual sampling in real time and underlines the importance of cross-modal predictions in adaptive visual behaviour and decision-making under uncertainty.

## Methods

### Participants

In a series of three experiments, we recruited 67 healthy human volunteers. Two participants were unable to complete the experiments due to nausea, and data from 2 more participants were excluded because do low data quality. Our analyses are based on data from the remaining 63 participants (N = 22, 21 and 20 in Experiments 1-3, respectively). On average our participants were 22 years old (Exp. 1: mean age = 20.8, range = 18-35, 18 women, all right-handed; Exp. 2: mean age = 22.9, range = 18-45, 11 women, 2 left-handed; Exp 3: mean age = 22.5, range = 18-55, 14 women, all right-handed). The sample size was defined based on prior studies using similar methods (Draschkow et al., 2022). All participants had normal or corrected-to-normal vision. All volunteers received financial compensation (30 PLN per hour). The study was approved by the institutional review board at the Jagiellonian University, and all participants gave written informed consent.

### Apparatus and virtual environment

Participants wore an HTC Vive Pro VR headset equipped with an integrated binocular eye tracker from Pupil Labs, which has an accuracy of approximately 0.5° of visual angle. The head-mounted display (HMD) featured two screens with a resolution of 1080×1200 pixels each, a refresh rate of 90 Hz, and a field of view of 100° horizontally × 110° vertically. Gaze position in 3D space was determined by intersecting the gaze vector with objects in the environment, as processed by Pupil Labs software^1^. Gaze data were sampled at the rate of 50 Hz. The headset’s position was tracked using two Lighthouse base stations, which emitted 60 infrared pulses per second, detected by 37 infrared sensors embedded in the HMD. Additional tracking was enhanced by an onboard accelerometer and gyroscope, allowing precise head rotation recordings with an accuracy of less than 0.001°. Participants sat on a chair and controlled the virtual bike using a gamepad, managing acceleration, deceleration, and steering from a first-person cyclist perspective.

The virtual environment was designed and rendered using the Unity Game Engine (Version 2023.9) on a Windows desktop computer, while eye-tracking data were recorded separately on a dedicated PC using Pupil Labs recording software. Within the VR environment, participants navigated a virtual urban setting as cyclists. The city was designed to be visually simple, featuring symmetrical buildings on both sides of the street, evenly distributed trees, sidewalks, and urban light poles. This minimized visuospatial complexity while maintaining a naturalistic, immersive experience. We designed an environment with low overall complexity by ensuring consistent street width (two designated bike lanes), symmetrical facades (in terms of height and architectural features), minimal clutter (repetitive street objects and trees), and limited pedestrian movement (based on Kondyli et al., 2020, 2023) Participants cycled along a dedicated cycling street, which had two lanes—one for the participant’s direction on the right and another for oncoming cyclists on the left. The route spanned 25 km, included 11 turns (both right and left), and incorporated various everyday traffic events such as traffic lights, bridge crossings, street obstacles, children playing, and pedestrian crossings.

### Procedure and tasks

Upon arrival, participants were familiarized with the head-mounted display (HMD) and the gamepad in a trial session within a virtual environment, where they cycled freely on bike lanes of a virtual city. Eye-tracking 9-point calibration was conducted before the start of the test session. All three experiments followed the same basic procedure and instructions. After completing the short practice session, participants were instructed to cycle toward a designated destination named “Liberty”, following the available traffic signs along the route. All participants started from the same location in the virtual city and followed an identical path toward the destination, where they encountered a finish line. Throughout the route, participants experienced “overtakes”, events in which an avatar vehicle (either a bicycle or a motorcycle) overtook the participant’s bike from the left or right side (Figure 1A-B). A typical overtaking event could last from 2.1 and 19.11 seconds (average 6 seconds) depending on the speed of the avatar and the participant. Participants were not given specific instructions on how to respond to these overtaking events to encourage natural reactions based on their everyday cycling experience. They could control both the direction and speed of their bike, with a maximum speed limit of 10 km/h. This speed constraint ensured that participants encountered everyday interaction events (e.g., pedestrian crossings, traffic lights) and the designed overtaking events within comparable time windows. Although participants had unlimited time to complete the navigation-cycling task, all of them finished the route within 15–25 minutes. Following the VR session, participants completed a questionnaire covering general demographic information (age, sex, dominant hand), VR experience assessment (perceived task difficulty, nausea), and previous experience with driving, cycling, and VR gaming (data not used in the current study).

### Experiment 1

In Experiment 1, participants encountered 40 instances of overtaking events along their cycling route. The number of overtakes were counterbalanced with 20 avatars overtaking the participant from the right side and 20 overtaking them from the left side. Because we had two types of overtaking avatars (i.e., motorbike and cyclist), we also counterbalanced the type of overtaking avatar. Specifically, we designed 10 overtaking events for each category: cyclist-left, cyclist-right, motorcyclist-left, and motorcyclist-right.

A 3D spatialized sound effect was implemented in Unity 3D to simulate natural overtaking incidents with a high degree of auditory realism. The sound was attached to the overtaking vehicle (e.g. bike, motorbike) as a dynamic audio source, leveraging Unity’s spatial audio system to enable precise localization in stereo headphones. The sound of a cyclist was represented by a typical bicycle recording, featuring the rhythmic whirring and clicking of the chain and gear sourced from an online media repository^2^. The motorcyclist’s sound included a low-frequency engine rumble and occasional revving. To approximate real-world conditions, the setup modeled natural sound propagation, including intensity increases as the source approached and a Doppler-like shift as the avatar passed and moved away.

The overtakes began with the “audio onset” where the sound originated from behind the participant’s bike at a distance of 20 meters, and increased in volume as the vehicle approached, culminating in the “Visual onset” when the vehicle became visible to the participant. In Unity’s 3D environment, audio lateralization is achieved by simulating spatial hearing based on the position of the sound source relative to the listener. As a sound moves left or right in 3D space, Unity automatically adjusts stereo panning and volume to reflect its direction and distance. This effect is controlled by the spatial blend setting, which determines how much the sound is influenced by 3D positioning versus being played equally in both ears. Concerning the settings on the “Audio Source” function in Unity, the spatializer was configured to calculate binaural cues for direction and distance, while the Doppler Effect dynamically altered the pitch based on the relative velocity of the vehicle. A logarithmic roll-off ensured natural attenuation of sound over distance, and environmental acoustics were enhanced using Unity’s reverb and occlusion settings where applicable. The Stereo Pan configuration was adjusted based on the overtaking side (right or left) to emphasize the corresponding headphone as the primary audio source. These combined settings created an immersive and realistic auditory experience for participants. No other background sounds were included. Overtaking instances were randomized along the route, with the time between events varying between 1.5 and 55 seconds. The overtaking bicycles traveled at a speed of 12 km/h, while the motorbikes moved at 15 km/h. The overtaking process lasted approximately 12 seconds for bicycles and 10 seconds for motorbikes. The avatar-cyclists and -motorcyclists varied in appearance, including differences in bike type, helmets, clothing, and hairstyles, and they were balanced for the left- and right-side overtakes.

### Experiment 2

This experiment followed the same general structure and used the same timing as Experiment 1, with one exception. Here, we introduced two types of overtaking events – congruent and incongruent events. In the *incongruent condition*, the sound source indicated that the avatar was overtaking from one side (e.g., left side), while the visual representation of the avatar entered into participant’s field of view from the opposite side (i.e., right side). The auditory stimuli were the same with the same timing, lateralization and the same Doppler effect. In the *congruent condition* the sound source indicated that the avatar was overtaking from one side (e.g., left side), and the visual representation of the avatar entered into participant’s field of view from the same side (i.e., left side). Participants encountered 40 overtaking events, with 30% of events being in the incongruent condition and 70% in the congruent condition.

### Experiment 3

In this experiment we maintained the general structure and timing of Experiment 1, with one key modification. Participants navigated a virtual city and encountered overtaking events similar to those in Experiments 1 and 2. From the total of 40 overtaking avatars 30% dropped the box right in front of the participant’s bike (12 events (Figure 2A), while 70% either carried a box and did not drop it (50% of cases, 14 events) or did not carry a box at all (50% of cases, 14 events). To examine the impact of auditory cues and their role in guiding adaptive behaviour, Experiment 3 required participants to navigate *with* predictions (i.e., audio-on condition) and *without* predictions (i.e., audio-off condition). Participants were instructed to cycle through the city as in Experiments 1 and 2, but they were also required to avoid colliding with the dropped boxes, necessitating timely reactions. Crucially, **time pressure** was introduced to test whether participants will be able to use the audio-based predictions to better anticipate overtaking events and prevent the collisions. Specifically, participants had approximately 500-800 ms between the moment the avatar became visible (visual onset) and the box drop. Failure to steer away from the box led to a collision after 700-800 ms. Each participant performed both conditions. The order of conditions was counterbalanced across participants to control for potential order effects. Half of the participants started with the audio-on condition, followed by the audio-off condition, while the other half experienced the reverse order. In both conditions, participants followed the same route. Key parameters—such as timing, the number of overtaking avatars, and the proportion of avatars carrying and dropping boxes—were kept consistent across conditions.

### Data preprocessing

The gaze data were recorded continuously at 50 Hz throughout the experimental sessions and analyzed in their unprocessed form, without being categorized into fixations and saccades. To ensure accuracy, only gaze points with confidence levels exceeding 70% were included in the analysis (Josupeit, 2023). Briefly, each gaze sample was associated with a confidence rate provided by the Pupil Labs detection algorithm, representing the estimated reliability of pupil detection on a scale from 0 (no confidence) to 100 (perfect confidence). Following Pupil Labs’ recommendations, data points with confidence values below 60% were considered unreliable and excluded from analysis. For more accuracy we decided to adjust the threshold higher at 65%. Horizontal gaze positions primarily ranged within 0.5 meters to the left or right of the central starting position. Less than 1% of the data fell outside this range, representing head movements rather than the subtle gaze shifts relevant to this study. These outliers were excluded to maintain the validity of the analysis and prioritize the fine-grained dynamics of gaze behaviour, which is central to the study’s focus on subtle gaze shifts. Since the study did not aim to investigate audio cues’ effects on head movements—owing to significant individual variability—data reflecting such movements were classified as outliers.

### Data temporal segmentation

To analyze gaze behaviour, all data were aligned relative to the moment the overtaking avatar entered the participant’s field of view, referred to as the “visual onset”, rather than the onset of the audio cue. This alignment was chosen for three reasons: 1) We expected shifts to be just in time or before the time when the avatar was entering into the visual field. 2) Early after audio onset the audio cues were not lateralized (no difference between left and right audio channel) to create a naturalistic impression that the avatar is approaching behind. Depending on the speed of the avatar and participant the cues started to be lateralized around 2 seconds after the audio onset. Consequently, we expected no gaze orienting early after auditory cue onset. 3) The duration of the entire overtaking event (i.e., the interval between audio onset and avatar entering the field of view) varied across trials because participants were free to navigate the virtual reality environment—adjusting their bike velocity, head movements, and path—resulting in differences in overall task duration. Eye-tracking data were segmented into three-second epochs around the visual onset, including two seconds before and one second after stimulus appearance. These segments were then averaged (Figure 1C). For the analyses presented in Figure 1D, 2C, and 3C, we additionally calculated the towardness index.

### Towardness index

To quantify gaze orienting towards auditory-gaze cued location we combined trials in which participant was overtaken from the right- and left-side into a single “towardness” index (for a similar approach see Draschkow et al., 2022). For each participant we first averaged all trials in which the participant was overtaken from both sides. Next, values for left and right overtakes were combined into a single metric by subtracting the left-side values from the right-side values after removing their respective baselines. Baseline was defined as the average gaze position of the first 25 samples within the time window from –2 to –1.5 seconds before the overtaking avatar entered the participants’ field of view (visual onset). This is a rather conservative approach because the auditory cue was typically audible for more than 2 seconds before the avatar entered the field of view. Control analyses using alternative baseline windows yielded consistent results, supporting the robustness of this approach.

### Statistics

Unless otherwise stated we used non-parametric tests (Wilcoxon signed-rank test, implemented via the Wilcoxon function from the scipy.stats module in Python). To quantify deviation of towardness index from the baseline, we compared each time point to the baseline interval. To control for multiple comparisons, we used Benjamini–Hochberg procedure. Note that we smoothed the towardness index for visualization purposes only (5-sample, 100 ms moving average) to reduce noise and enhance clarity. Comparisons between conditions—such as overall duration of gaze fixation on the avatar between congruent and incongruent sound conditions in Experiment 2, and average reaction time as well as collision counts between audio-on and audio-off conditions in Experiment 3—were also conducted using the Wilcoxon signed rank test. These data were visualized using box plots with individual participant’s data overlaid.

### Area of interest (AOI) analysis

Concerning the analysis of gaze within areas of interest (AOIs), in our studies these areas were defined as the three-dimensional spaces encapsulating virtual avatars and their vehicles, such as the bikes or motorbikes, that overtook the participant. AOIs were implemented as 3D colliders within the VR environment to track gaze intersection with these moving objects. Dwell time within AOIs was analyzed for the first second after visual onset, as this period was most relevant for detecting gaze bias. Gaze hits beyond this time window were excluded, as object visibility varied due to avatar velocity, environmental geometry, and participant behaviour, such as voluntary head movements. For comparisons involving the congruent and incongruent conditions (Experiment 2), a transparent three-dimensional object was created to represent the audio source in Unity. This object, anchored to the audio source, follows a trajectory that mirrors the path and volume of the overtaking avatar relative to the participant’s trajectory. The volumes surrounding the audio source and the visual avatar were defined as areas of interest (AOIs). Gaze hits within these AOIs were tracked to analyze participants’ gaze behaviour in relation to the spatial and auditory characteristics of the scenario.

## Acknowledgments

This research was funded by a grant from National Science Center of Poland (OPUS 23: 2022/45/B/HS6/04097).

## Data and Code availability

Video material of example trials from Experiments 1, 2, and 3, data and code are available at https://osf.io/4hjwf/

## Footnotes

1 Details about the Pupil Labs software are available at https://docs.pupil-labs.com/core/software/pupil-player/

2 We used the bicycle-wheel-spinning-49716.mp3 by the free-sound community, https://pixabay.com/sound-effects/

